# Evidence for host-dependent RNA editing in the transcriptome of SARS-CoV-2

**DOI:** 10.1101/2020.03.02.973255

**Authors:** Salvatore Di Giorgio, Filippo Martignano, Maria Gabriella Torcia, Giorgio Mattiuz, Silvestro G. Conticello

**Author notes:** These authors contributed equally (Salvatore Di Giorgio, Filippo Martignano, Giorgio Mattiuz, Silvestro G. Conticello).

## Abstract

The COVID-19 outbreak has become a global health risk and understanding the response of the host to the SARS-CoV-2 virus will help to contrast the disease. Editing by host deaminases is an innate restriction process to counter viruses, and it is not yet known whether it operates against Coronaviruses. Here we analyze RNA sequences from bronchoalveolar lavage fluids derived from infected patients. We identify nucleotide changes that may be signatures of RNA editing: Adenosine-to-Inosine changes from ADAR deaminases and Cytosine-to-Uracil changes from APOBEC ones. A mutational analysis of genomes from different strains of human-hosted *Coronaviridae* reveals mutational patterns compatible to those observed in the transcriptomic data. Our results thus suggest that both APOBECs and ADARs are involved in Coronavirus genome editing, a process that may shape the fate of both virus and patient.

**For the casual Reader:** Just to make a few things clear:

- RNA editing and DNA editing are PHYSIOLOGICAL processes. Organisms uses them to (a) try to fight viruses, (b) increase heterogeneity inside cells (on many levels), (c) recognise their own RNA.
- our work suggests that: (a) cells use RNA editing in trying to deal with Coronaviruses. We don't know to what extent they succeed (and it would be nice if we could help them). (b) Whatever happens, mutations inserted by RNA editing fuel viral evolution. We don't know whether viruses actively exploit this.
- If you (scientist or not) think our work suggests ANYTHING ELSE, contact us. It can be a first step to help fight these !@#$ coronavirus, or towards a Nobel prize - but we need to discuss it thoroughly.
- If you think these cellular processes are fascinating, join the club and contact us. We can have a nice cup of tea while chatting how wondrous nature is at coming up with extraordinary solutions…

## Introduction

Emerging viral infections represent a threat to global health, and the recent outbreak of Novel Coronavirus Disease 2019 (COVID-19) caused by Severe Acute Respiratory Syndrome Coronavirus 2 (SARS-CoV-2, Novel Coronavirus, 2019-nCoV) exemplifies the risks (*1*, *2*). As viruses are obligate intracellular parasites, organisms have evolved innate immune processes to sense and counter the viruses. Among these, RNA and DNA editing mediated by endogenous deaminases can provide a potent restriction against specific viruses. Two deaminase families are present in mammalian species: the ADARs target double stranded RNA (dsRNA) for deamination of Adenines into Inosines (A-to-I) (*3*, *4*), and the APOBECs deaminate Cytosines into Uracils (C-to-U) on single-stranded nucleic acids (ssDNA and ssRNA) (*5*, *6*). During viral infections ADARs act either directly, through hypermutation of the viral RNA, or indirectly, through editing of host transcripts that modulate the cellular response (*7–18*). On the other hand, APOBECs target the viral genome, typically DNA intermediates (*19–26*), either through C-to-U hypermutation or through a non-enzymatic path that interferes with reverse transcription (*27*, *28*). Some APOBEC3 proteins can interfere *in vitro* with *Coronaviridae* replication, yet it is not clear whether their enzymatic activity is involved (*29*). Eventually though, these restriction systems can also be exploited by the viruses to support their infectivity and increase their evolutionary potential (9, *11–15*, *30–32*).

## Results

To assess whether RNA editing could be involved in the response to SARS-CoV-2 infections, we started from publicly available RNA sequencing datasets from bronchoalveolar lavage fluids (BALF) obtained from patients diagnosed with COVID-19. While transcriptomic data for all samples could be aligned to the SARS-CoV-2 reference genome, the quality of the sequencing varied and only eight samples had coverage and error rates suitable for the identification of potentially edited sites (**Data S1**). We called single nucleotide variants (SNVs) on these eight samples (*33*, *34*) using REDItools 2 (*35–37*) and JACUSA (*38*) using the following thresholds: reads supporting the SNV ≥4, allelic fraction ≥0.5%, coverage ≥20, quality of the reads >25, base quality >35 (**Fig. S1A**). The two pipelines gave comparable results with ~50% of the SNV positions called by both (**Fig. S1B, Fig S2**). We identified 910 SNVs common to REDItools 2 and JACUSA, ranging from 24 to 238 SNVs per sample (**Fig. 1, Data S3**). Given the thresholds used to call the SNV, samples with lower sequencing depths displayed lower numbers of SNVs.

**Fig. 1.**
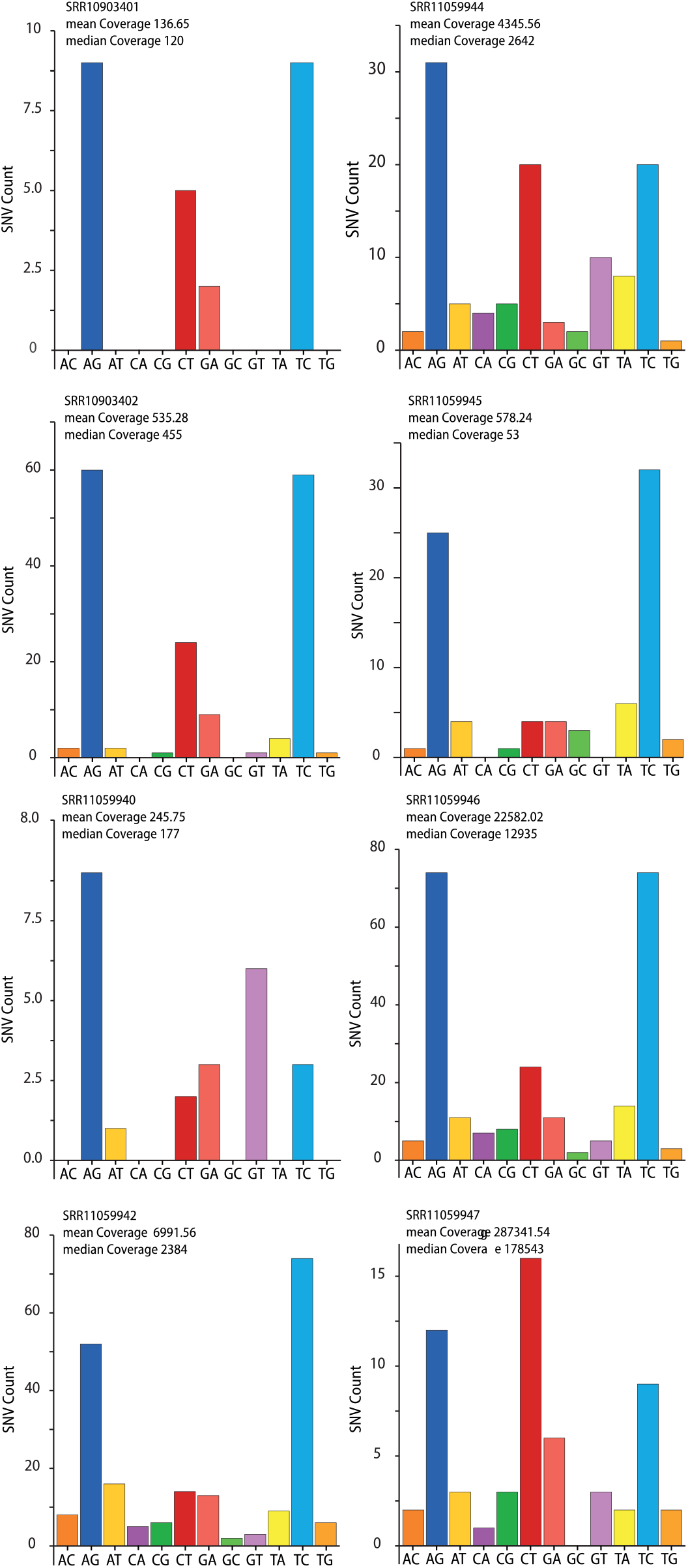
Single-nucleotide variants (SNV) identified in SARS-CoV-2 transcriptomes. The bar charts show the number of SNVs identified in each SARS-CoV-2 transcriptome for each SNV type (e.g. A>C, AC). The sequencing depth for each sample is indicated.

While the weight of each SNV type varies across samples (**Fig. 1**), a bias towards transitions is always present, which is even more evident when all mutational data are pooled (**Fig. 2A, B**). This holds true even when only SNVs recurring in more samples are considered (**Fig. 2C**).

**Fig. 2.**
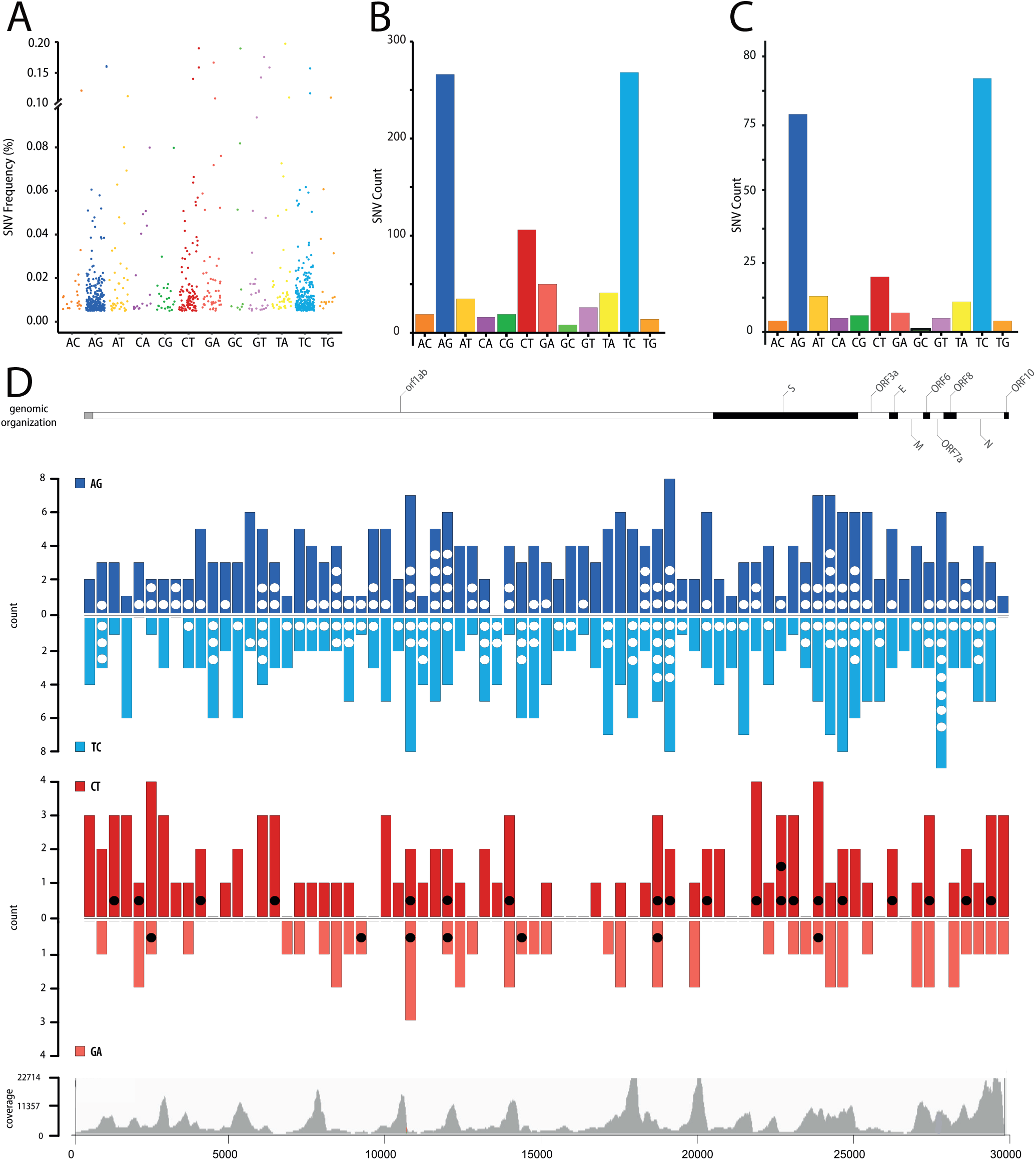
Single-nucleotide variants (SNV) identified in SARS-CoV-2 transcriptomes. (**A**) Allelic fraction and (**B**) number of SNVs for each nucleotide change in the entire dataset and (**C)** for SNVs recurring in at least two samples. **(D)** Distribution of SNVs across the SARS-CoV-2 genome. A-to-G (*blue*) and C-to-U (*red*) SNVs are grouped in 400nt bins and plotted above (AG and CT) or below the line (TC and GA) based on the edited strand. Dots (*white/black*) indicate recurring SNVs. Genetic organization of SARS-CoV-2 (top), the *dark*/*white* shading indicates the viral coding sequences; coverage distribution of all analyzed samples (bottom).

SNV frequency and number of transversions are compatible with mutation rates observed in Coronaviruses (10^−6/−7^, (*39*) and commonly associated to the RNA dependent RNA polymerases (RdRp). RdRp are error-prone and are considered the main source of mutations in RNA viruses. However, the Coronavirus nsp14-ExoN gene provides a form of error correction (*40*), which is probably the reason mutation rates in Coronaviruses are lower than those observed in RNA viruses with smaller genomes. The mutational spectrum in SARS quasispecies presents a very weak bias towards U-to-G. Inactivation of nsp14-ExoN error correction reveals the mutational spectrum of the RdRp, which is quite different from the pattern we observe (i.e. main changes are C-to-A, followed by U-to-C, G-to-U, A-to-C and U-to-G) (*41*). As such we would consider that SNVs deriving from RdRp errors represent a marginal fraction of the SNVs in the SARS-CoV-2 samples.

The bias towards transitions-mainly A>G/T>C changes-resembles the pattern of SNVs observed in human transcriptomes (*42*) or in viruses (*8*, *10*, *18*), where A>G changes derive from deamination of A-to-I mediated by the ADARs. It is thus likely that also in the case of SARS-CoV-2 these A>G/T>C changes are due to the action of the ADARs.

C>T and G>A SNVs are the second main group of changes and could derive from APOBEC-mediated C-to-U deamination. Contrary to A-to-I, C-to-U editing is a relatively rare phenomenon in the human transcriptome (*42*) and, with regard to viruses, it has been associated only to positive-sense ssRNA Rubella virus (*32*), where C>T changes represent the predominant SNV type. The observation that only A-to-I editing is present in RNA viruses that infect non-vertebrate animals, where RNA-targeting APOBECs are not present (*10*, *18*), supports the hypothesis that APOBECs are involved in the RNA editing of this human-targeting virus.

A third group of SNVs, A>T/T>A transversions, is also present. While this type of SNVs has been described (*43*), its origin is yet unknown.

A>G and T>C changes are evenly represented for SNV frequency (**Fig. 2A**), for number of unique SNVs (**Fig. 2B, C**), and for distribution on the viral genome (**Fig. 2D**). As ADARs target dsRNA, this suggests that dsRNA encompasses the entire genome. While dsRNA in human transcripts are often driven by inverted repeats, the most likely source of dsRNA in the viral transcripts is the replication, where both positive and negative strands are present and can result in wide regions of double-stranded RNA. Contrary to A-to-I changes, C-to-U changes are biased towards the positive-sense strand (**Fig. 2B, C, D,***p* < 0.0001). Since ADARs and APOBECs target selectively dsRNA and ssRNA, such distribution could derive from the presence at all times of RNA in a dynamic equilibrium between double-strandedness - when negative-sense RNA is being transcribed- and single-strandedness-when nascent RNA is released. Some areas seem to bear less SNVs, but this might be related to lower sequencing depth in those regions. As APOBEC deaminases preferentially target cytosines within specific sequence contexts, we analyzed the nucleotide context of A-to-I and C-to-U SNVs in the viral genome (**Fig. 3A**, **B**). A slight depletion of G bases in position −1 is present at A-to-I edited positions. The depletion is not as strong as the signal previously reported in human transcripts (*44–47*). The low editing frequencies we observe resembles the editing present on human transcripts containing Alu sequences, which were in a limited number in those early datasets. Indeed, there is no evidence of a sequence context preference if we use a larger dataset such as REDIportal (*48*), which includes >1.5 M sites in Alu repeats (**Fig. S3**).

**Fig. 3.**
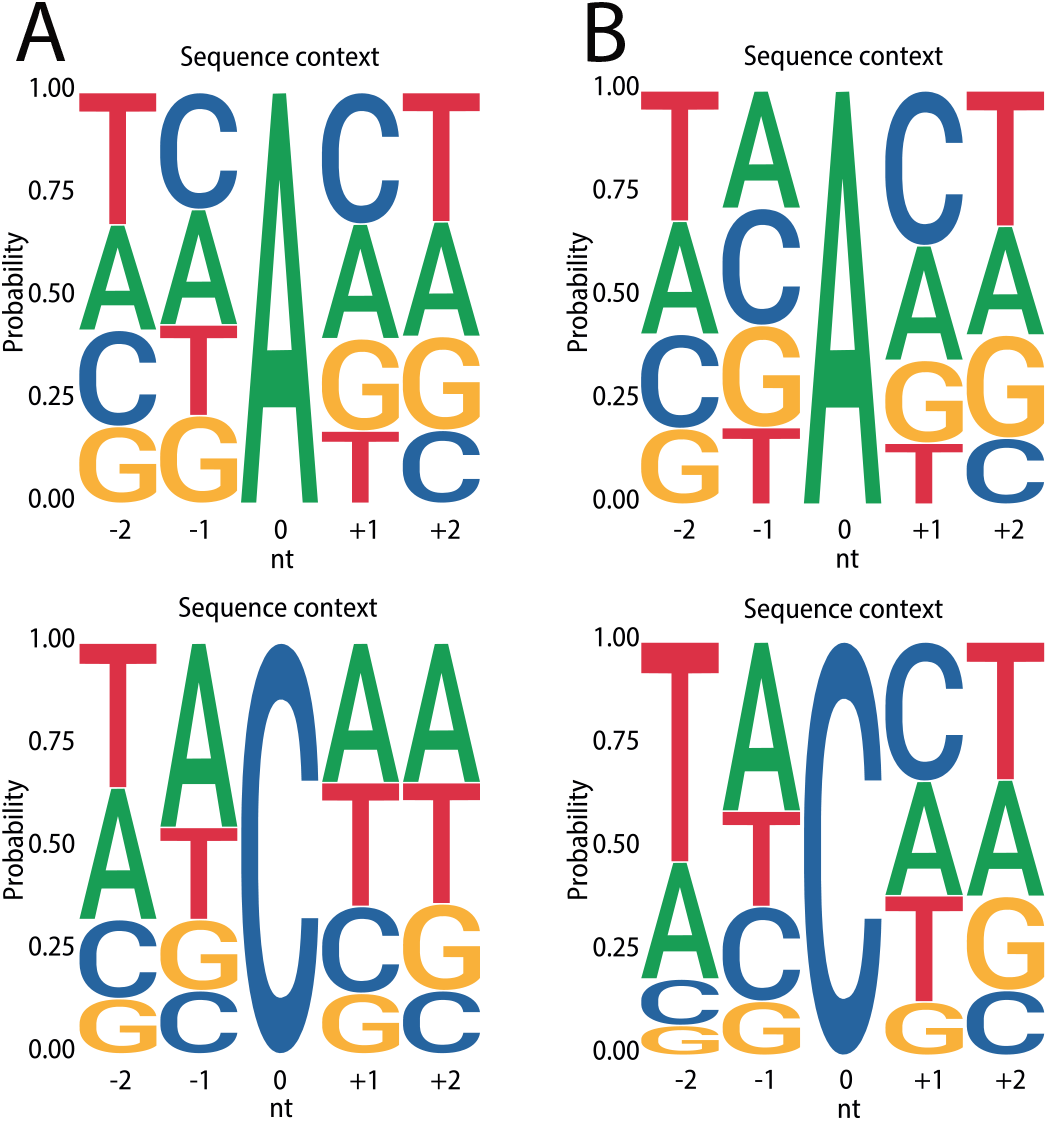
Sequence contexts for SARS-CoV-2 RNA edited sites. **(A)** Local sequence context for A-to-I and C-to-U edited sites in the viral transcriptome and (**B**) for recurring sites.

On the other hand, C-to-U changes preferentially occur 3’ to thymines and adenines, a sequence context that resembles the one observed for APOBEC1-mediated deamination ([AU]C[AU]) (*49*, *50*).

We then aligned available genomes from SARS-CoV-2, Middle-East Respiratory Syndrome-related Coronavirus (MERS-CoV), and the Severe Acute Respiratory Syndrome Coronavirus (SARS-CoV) to test whether RNA editing could be responsible for some of the mutations acquired through evolution. Indeed, the genomic alignments reveal that a substantial fraction of the mutations in all strains could derive from enzymatic deaminations (**Fig. 4A, B, C**), with a prevalence of C-to-U mutations, and a sequence context compatible with APOBEC-mediated editing exists also in the genomic C-to-U SNVs (**Fig. 4D, E, F**).

**Fig. 4.**
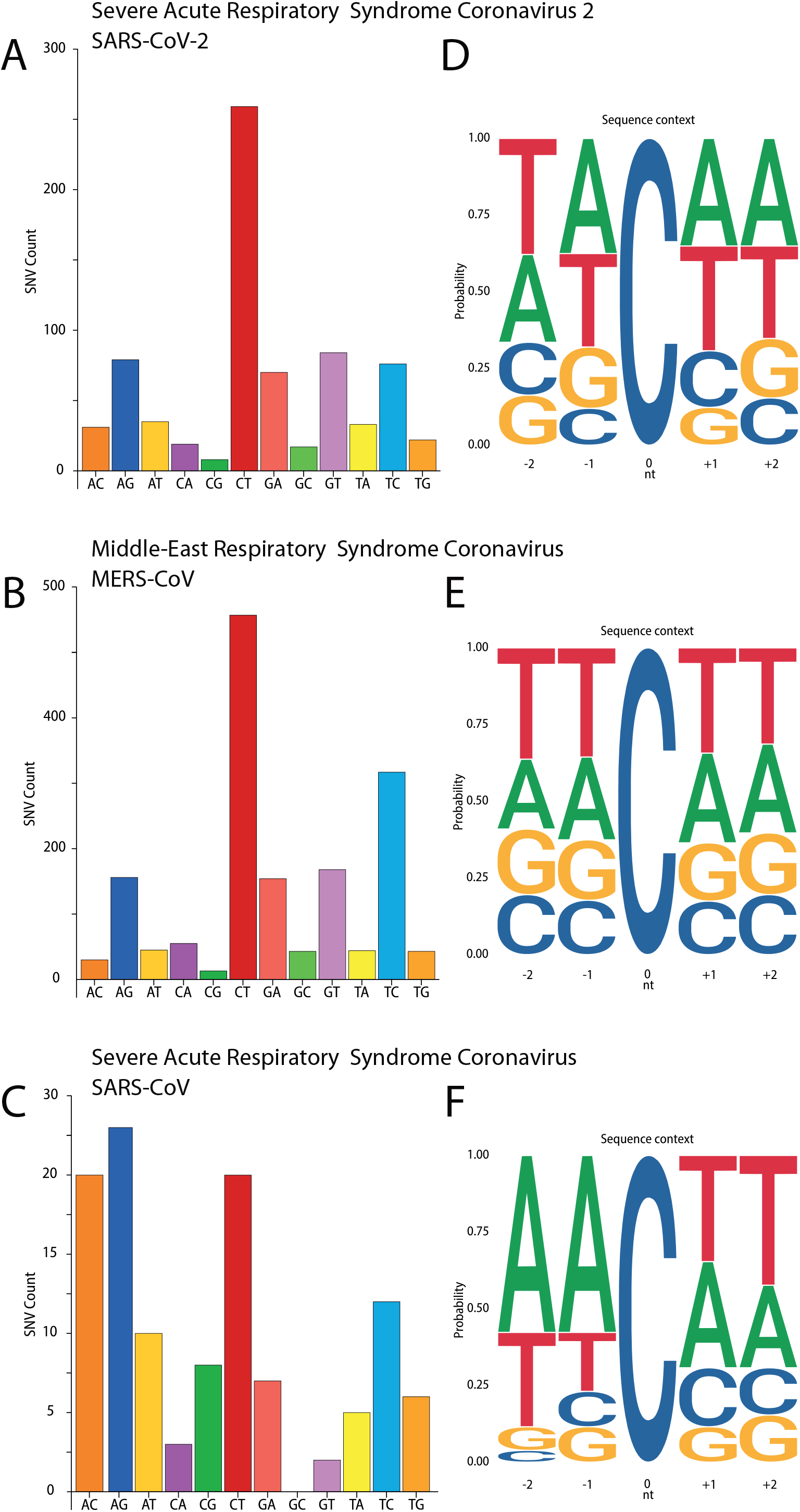
Nucleotide changes across *Coronaviridae* strains. **(A, B, C)** Number of SNVs for each nucleotide change and (**D, E, F**) local sequence context for C-to-U edited sites in genome alignments from SARS-CoV-2 **(A, D)**, human-hosted MERS-CoV **(B, E)**, and human-hosted SARS-CoV **(C, F)**.

## Discussion

Considering the source of our analysis-metagenomic sequencing-, we wonder whether the low-level editing we observe (~1%) reflects the levels of editing of the viral transcripts inside the cells. Indeed, beside a small fraction of cellular transcripts edited at high-frequency, most ADAR-edited sites in the human transcriptome (typically inside Alu sequences) present editing levels of ~1% (*4*, *42*, *51*). It has been shown that a fraction of the cellular transcripts are hyperedited by ADARs (*52–54*). While we were unable to observe hyperedited reads in the metagenomic samples, it is possible that hyperedited transcripts fail to be packaged into the virus.

With regard to APOBEC-mediated RNA editing, its detection in the viral transcriptomes is already indicative, as this type of editing is almost undetectable in human tissues (*42*). Such enrichment points either towards an induction of the APOBECs triggered by the infection, or to specific targeting of the APOBECs onto the viral transcripts. The APOBECs have been proved effective against many viral species in experimental conditions, yet, until now their mutational activity in clinical settings has been shown only in a handful of viral infections (*19–26*) through DNA editing- and in Rubella virus, on RNA (*32*).

As in Rubella virus, we observe a bias in APOBEC editing towards the positive-sense strand. This bias and the low editing frequencies might be indicative of the dynamics of the virus, from transcription to selection of viable genomes. It is reasonable to assume that sites edited on the negative-sense strand will result in a mid-level editing frequency, as not all negative-sense transcripts will be edited (**Fig. 5A**). On the other hand, editing of the positive-sense strand can occur upon entry of the viral genome, thus yielding high-frequency editing (**Fig. 5B**), or after viral genome replication, resulting in low-frequency editing (**Fig. 5C**). Indeed, lack of a sizable fraction of highly edited C>T SNVs suggests that APOBEC editing occurs late in the viral life-cycle (**Fig. 5C**). Yet, since they occur earlier, G>A SNVs should be closer in number to C>T ones and with higher levels of editing, which is not what we observe (**Fig. 2A, B, C**). The overrepresentation of C>T SNVs could be due to an imbalance towards positive-sense transcripts, as these are continuously generated from the negative-sense ones (and double-stranded hybrid RNAs are lost). However, the editing frequencies of G>A SNVs should be much higher as G>A SNVs are generated upstream to the C>T ones. A more fitting explanation is that editing of the negative-sense transcripts results in loss of the edited transcript (**Fig. 5D**), possibly because editing triggers nonsense mediated decay (*55*), thus lowering the chances of the edited site to be transmitted.

**Fig. 5.**
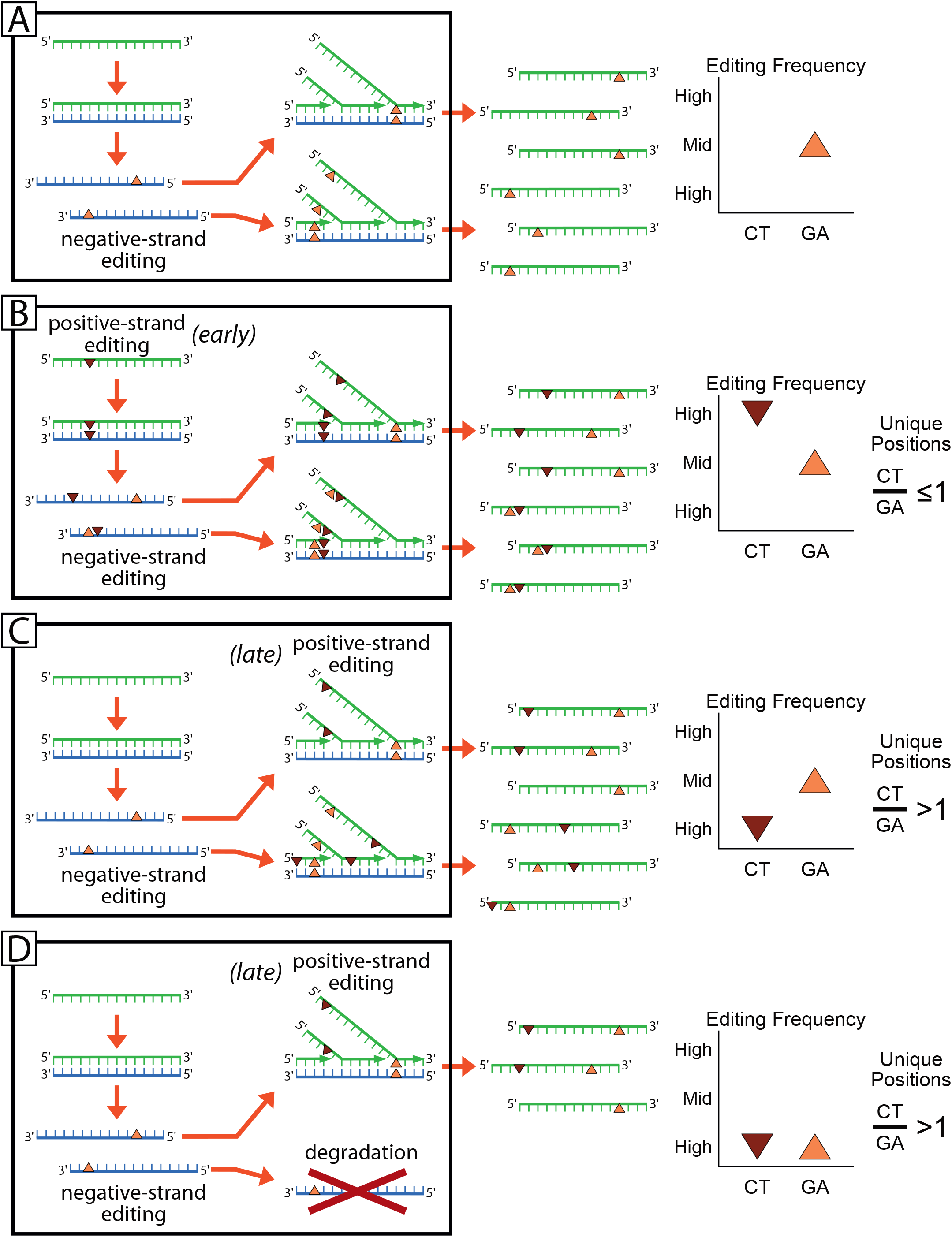
Model of APOBEC RNA editing on SARS-CoV-2 transcriptome. The 4 panels model the editing frequencies and the C>U/G/A ratios expected from 4 different scenarios: (A) C-to-U editing on the negative-sense transcripts; (B) ‘early’ editing on the viral genomes before viral replication; (C) ‘late’ editing after viral replication; (D) ‘late’ editing after viral replication with loss of negative-sense transcripts. Red dots indicate editing on the positive-sense transcript; Orange dots indicate editing on the positive-sense transcript. Green and Blue segments indicate positive- and negative-sense viral transcripts, respectively.

Since most of the APOBECs are unable to target RNA, the only well characterized cytidine-targeting deaminase is APOBEC1, mainly expressed in the gastrointestinal tract, and APOBEC3A (*56*), whose physiological role is not clear. As with A-to-I editing, it will be important to assess the true extent of APOBEC RNA editing in infected cells.

The functional meaning of RNA editing in SARS-CoV-2 is yet to be understood: in other contexts, editing of the viral genome determines its demise or fuels its evolution. For DNA viruses, the selection is indirect, as genomes evolve to reduce potentially harmful editable sites (e.g. (*18*), but for RNA viruses this pressure is even stronger, as RNA editing directly affects the genetic information and efficiently edited sites disappear.

A comparison of the SNV datasets from the transcriptomic and from the genomic analyses reveals a different weight of A-to-I and C-to-U changes (**Fig. 2B, Fig. 4A**), with an underrepresentation of A-to-I in the viral genomes. As our analysis underestimates the amount of editing due to strict parameters used, it could be possible that A-to-I SNVs are not fixed in the viral population when ADAR editing is such that impairs the viral genome.

An analysis of the outcomes of the mutations is difficult due to the low numbers of events collected so far, but there are some trends that might be suggestive (**Data S2**). C-to-U changes leading to Stop codons are overrepresented in the transcriptomic data but -as expected-disappear in the genomic dataset. This might point-again- to an antiviral role for these editing enzymes. There is also an underrepresentation of C>T missense mutations, but its meaning is difficult to interpret.

Finally, this analysis is a first step in understanding the involvement of RNA editing in viral replication, and it could lead to clinically relevant outcomes: (a) if these enzymes are relevant in the host response to Coronavirus infection, a deletion polymorphism quite common in the Chinese population, encompassing the end of APOBEC3A and most of APOBEC3B (*57*, *58*) could play a role in the spread of the infection. (b) Since RNA editing and selection act orthogonally in the evolution of the viruses, comparing genomic sites that are edited with those that are mutated could lead to the selection of viral regions potentially exploitable for therapeutic uses.

## Materials and Methods

### Sequencing Data

RNA sequencing data available from projects PRJNA601736, PRJNA603194 and PRJNA605907 were downloaded from NCBI (https://www.ncbi.nlm.nih.gov/sra/) using the FASTQ-dump utilities from the SRA-toolkit with the command line:

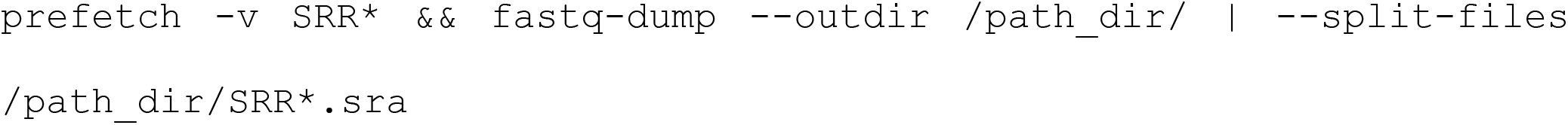

Since most of the reads of samples from PRJNA605907 were missing their mate, forward-reads and reverse-reads from these samples have been merged in a single FASTQ, which is treated as a single-end experiment. Details of the sequencing runs are summarized in **Data S1**.

### Data pre-processing

SRR11059940, SRR11059941, SRR11059942 and SRR11059945 showed a reduced quality of the sequencing in the terminal part of the reads.

We used TRIMMOMATIC (*59*) to trim the reads of those samples to 100 bp, with the following command line:

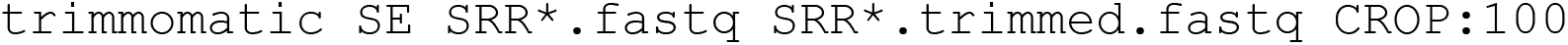

We aligned the FASTQ files using Burrows-Wheeler Aligner (BWA) (*60*) using the official sequence of SARS-CoV-2 (NC_045512. 2) as reference genome. After the alignments BAM files were sorted them using SAMtools (*61*).

Command line used for paired-end samples:

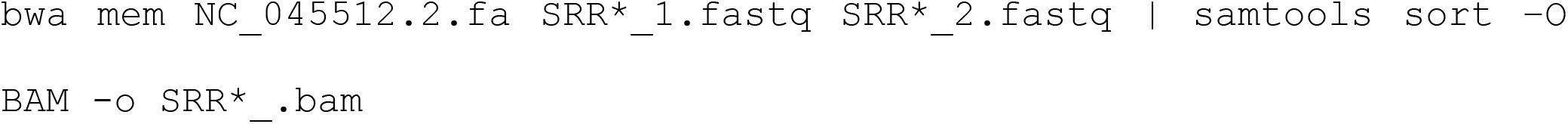

Command line used for single-end samples:

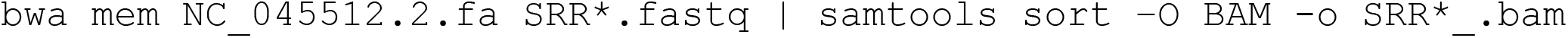

The aligned bams have been analysed with QUALIMAP (*62*). Due to a high error rate reported by QUALIMAP, samples SRR11059943 and SRR10971381 have been removed from the analysis.

### Single nucleotide variant (SNV) calling

A diagram of the entire pipeline is shown in **Fig. S1A**. We used RediTools 2 (*35*, *37*) and JACUSA (*38*) to call the SNVs using the command line:

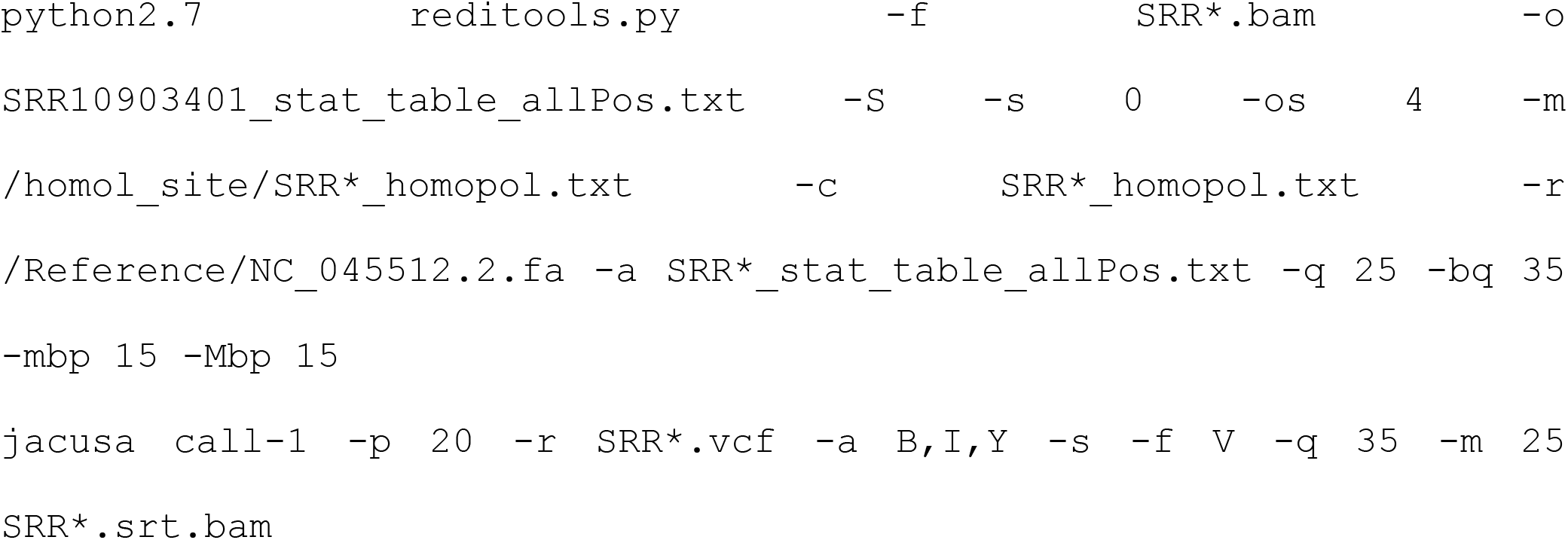

With regard to REDItools 2, we removed all SNVs within 15 nucleotides from the beginning or the end of the reads to avoid artifacts due to misalignments.

To avoid potential artifacts due to strand bias, we used the AS_StrandOddsRatio parameter calculated following GATK guidelines ((https://gatk.broadinstitute.org/hc/en-us/articles/360040507111-AS-StrandOddsRatio), and any mutation with a **AS_StrandOddsRatio** > 4 has been removed from the dataset.

Bcftools (*61*) has been used to calculate total allelic depths on the forward and reverse strand (ADF, ADR) for **AS_StrandOddsRatio** calculation, with the following command line:

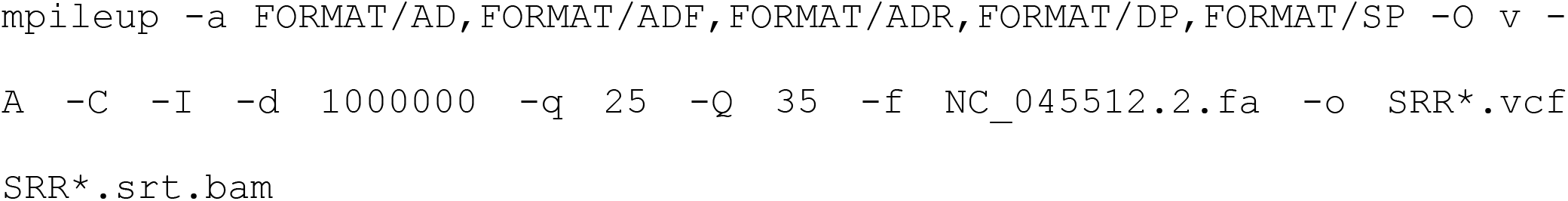

Mutations common to the datasets generated by Reditools 2 and JACUSA were considered (n = 910, **Fig. S2, Data S3**). The threshold we used to filter the SNVs is based on minimum coverage (20 reads), number of supporting reads (at least 4 mutated reads), allelic fraction (0.5%), quality of the mapped reads (> 25), and base quality (>35). In the dataset there were only 6 SNVs with allelic fractions in the range 30-85% (C>T, 1; T>C, 3; G>T, 2). Since there were no SNVs with higher allelic fractions, we presume that all samples originated from the same viral strain.

Recurring SNVs have been defined as the SNVs present in at least two samples. In order to overcome the problem of samples with lower sequencing depth, we used the positions of the SNVs common to both REDItools 2 and JACUSA to call again the SNVs irrespectively of the number of supporting reads.

### Data manipulation

R packages (Biostrings, rsamtools, ggseqlogo ggplot2, splitstackshape) and custom Perl scripts were used to handle the data.

### Sequence context analysis

Logo alignments were calculated using ggseqlogo, using either the pooled dataset of the dataset of recurring SNVs. Logo alignments the human edited sites were performed using ADAR sites from REDIportal (*48*) that were shared by at least 4 samples. SARS-CoV-2, SARS and MERS genomic data were prepared for the Logi alignment using the GenomicRanges R package (*63*).

### SNV calling in genomic data from SARS-CoV-2, SARS and MERS

The viral genomic sequences of MERS (taxid:1335626) and SARS (taxid:694009) were selected on NCBI Virus (https://www.ncbi.nlm.nih.gov/labs/virus/vssi/#/) using the query: Host : Homo Sapiens (human), taxid:9606;-Nucleotide Sequence Type: Complete. They were aligned using the “Align” utility. Consensus sequences of SARS and MERS genomes were built using the “cons” tool from the EMBOSS suite (http://bioinfo.nhri.org.tw/gui/) with default settings.

SARS-CoV-2 genomic sequences were downloaded from GISAID (https://www.gisaid.org/) and aligned with MUSCLE (*64*).

SNVs have been called with a custom R script, by comparing viral genome sequences to the respective consensus sequence or, for SARS-CoV-2, to the NC_045512.2 reference sequence. SNVs, viral consensus sequences, and *Coronoviradae* genome sequences identifiers, are provided in **Data S3**, **S4** and S5.

### SNVs annotation

SNVs (from both genomic and somatic SNVs sets) occurring on coding sequences have been annotated with custom R scripts to determine the outcome of the nucleotide change (nonsense/missense/synonymous mutation). A summary is reported in **Data S2**.

### Statistical Analysis

fisher.test() function from the R base package has been used for all the statistical tests. To test the significance of C-to-U bias on the positive strand, we compared C>T/G>A SNV counts to the count of C/G bases on the reference genome. For P-values of “RNA vs Reference”, “DNA vs Reference”, and “genome vs RNA” 2×2 contingency tables have been generated as shown in **Data S2**.

## Supplementary Materials

**Figs. S1 to S3**

**Data S1**: RNA sequencing details for each SARS-CoV-2 sample;

**Data_S2**: Missense/Nonsense/Synonymous mutations in SARS-CoV-2 transcriptomic and genomic data;

**Data S3**: List of SNVs on SARS-CoV-2 identified by both REDItool 2 and JACUSA; List of recurring SNVs.

**Data S4**: MERS and SARS consensus sequences obtained from deposited genomic sequences;

**Data S5**: accession numbers of the Coronavirus (SARS-CoV-2, MERS and SARS) genomic sequences used in the multi-alignments.

**Data S6**: SNVs in *Coronaviridae* genomes.

**Data S7**: List of all the authors, originating and submitting laboratories, whose sequences have been used.

## Acknowledgments

In memory of Li Wenliang, Carlo Urbani and of all the doctors and health workers who endangered their lives in the fight against epidemics. We acknowledge and thank all authors that are sharing their data. We gratefully acknowledge the authors, originating and submitting laboratories, of the sequences from GISAID’s EpiFlu Database on which this research is based. The list is detailed in Data file S6.

## Author contributions

Conceptualization: SDG, FM, MGT, GM, SGC; Formal Analysis, Investigation, Software: SDG, FM; Visualization: GM; Writing – original draft: GM and SGC; Writing – review&editing: SDG, FM, MGT, GM, SGC

## Data and materials availability

Sequencing and genomic data are available through NCBI SRA and Virus repositories, and GISAID.

## Supplementary Figures

**Fig. S1.**
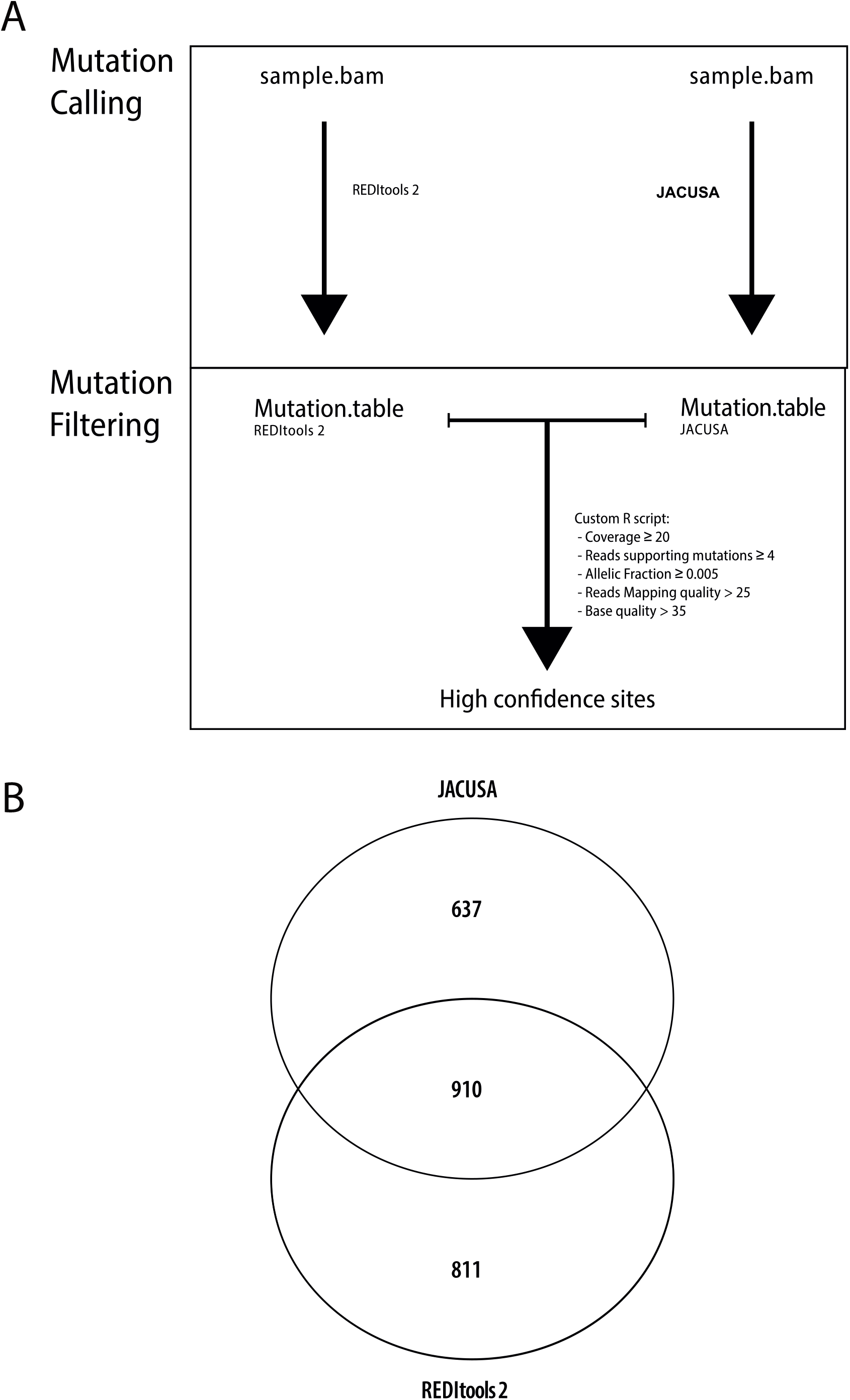
**(A)** Workflow: Mutation Calling and Mutation Filtering by REDItools 2 and JACUSA. **(B) Venn Diagram of the SNVs identified by REDItool 2 and JACUSA**.

**Fig. S2.**
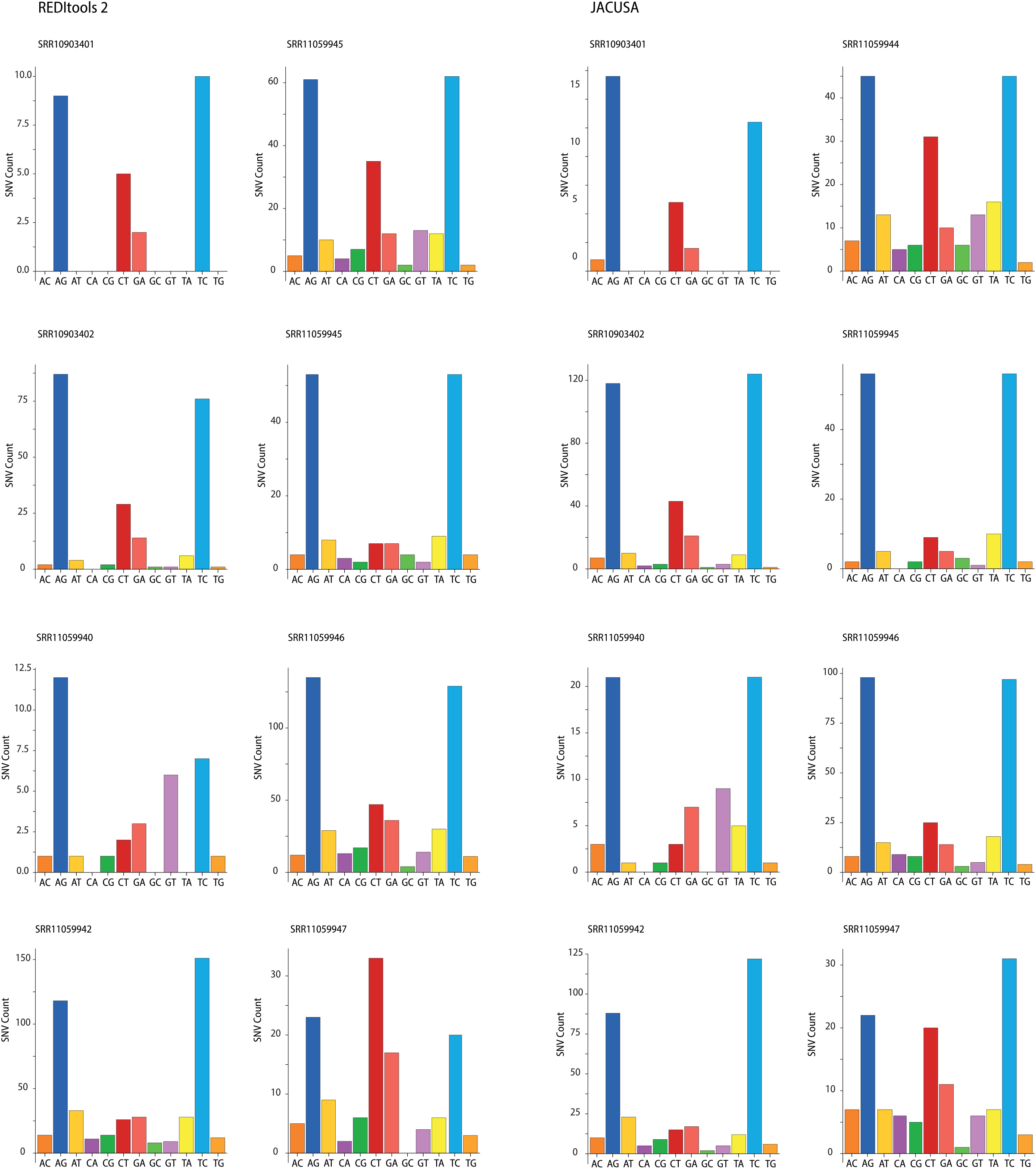
Single-nucleotide variants (SNV) identified in SARS-CoV-2 transcriptomes by REDItools 2 (left) and JACUSA (right). The bar charts show the number of SNVs for each 2019-nCoV transcriptome (e.g. A>C, AC).

**Fig S3.**
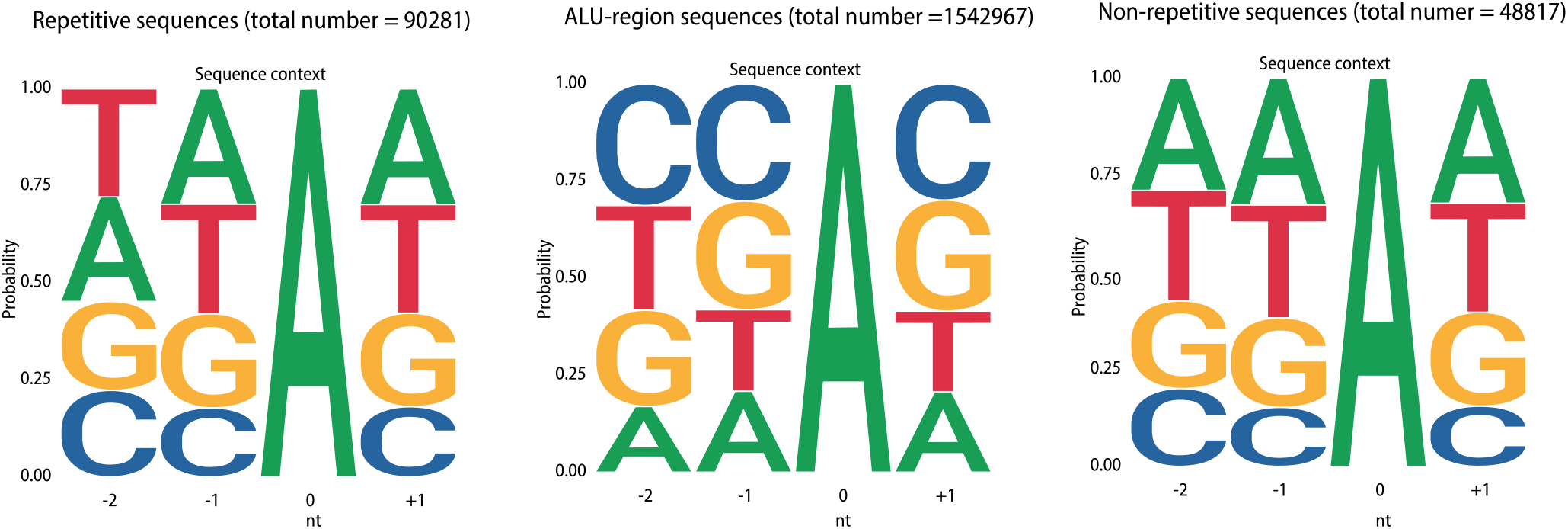
A-to-I mutation context in human transcriptome. Local sequence context for A-to-I sites calculated using the dataset from REDIportal.

